# pamlr: a toolbox for analysing animal behaviour using pressure, acceleration, temperature, magnetic and light data in R

**DOI:** 10.1101/2021.08.02.454456

**Authors:** Kiran L. Dhanjal-Adams, Astrid S. T. Willener, Felix Liechti

## Abstract

Light-level geolocators have revolutionised the study of animal behaviour. However, lacking precision, they cannot be used to infer behaviour beyond large-scale movements. Recent technological developments have allowed the integration of barometers, magnetometers, accelerometers and thermometers into geolocator tags, offering new insights into the behaviour of species which were previously impossible to tag.
Here, we introduce an R toolbox for identifying behavioural patterns from multisensor geolocator tags, with functions specifically designed for data visualisation, calibration, classification and error estimation. Some functions are also tailored for identifying specific behavioural patterns in birds (most common geolocators-tagged species), but are flexible for other applications.
Finally, we highlight opportunities for applying this toolbox to other species beyond birds, the behaviours they might identify and their potential applications beyond behavioural analyses.

**Data archiving:** Currently, the package is on github and will be submitted to CRAN after review. The supporting code (package manual) for this paper is also on https://kiranlda.github.io/PAMLrManual/index.html, but will later be hosted by the Swiss Ornithological Institute.

**Summary:** pamlr: a toolbox for analysing animal behaviour using pressure, acceleration, temperature, magnetic and light data in R

## 1 Introduction

Light-level geolocators (GLS) have revolutionised the study of animal behaviour. Thanks to their relatively low cost and light weight (<1g), they have facilitated the study of species which would otherwise be impossible to track. However, GLS tags rely on the use of light measurements to calculate an animal’s location (Frisius, 1544; Shaffer et al., 2005), creating a spatial accuracy of tens or even hundreds of kilometres (Simeon Lisovski et al., 2012), and thus restricting behavioural analyses to finding periods when the animal has performed movements larger than the error range of the device – typically migration.

To overcome this limitation, geolocators have recently been integrated with barometers, accelerometers, thermometers and magnetometers, creating opportunities for studying previously unobserved behaviours in small airborne birds, but also bats. Indeed, barometers allow us to explore an bird’s behaviour in a third dimension – height and depth – and can inform on flying, flocking, diving and foraging behaviours (Dhanjal-Adams et al., 2018; Dreelin, Shipley, & Winkler, 2018; Meier et al., 2018; Sjöberg et al., 2018). Accelerometers can be used to understand resting behaviour, migration timing, and how long animals remain airborne (Liechti, Witvliet, Weber, & Bächler, 2013; Hedenström et al., 2016) and to identify endurance flights (Liechti et al., 2018; Sjöberg et al., 2018). Thermometers can inform on habitat usage (Shaffer et al., 2005; Edwards, Quinn, & Thompson, 2016), and magnetometers can be used to understand bearing and direction (Bidder et al., 2015).

Though all of these sensors have previously been integrated into GPS tags with an increasing number of methods available for identifying behavioural states, GPS analyses rely heavily (i) on more precise location estimates to infer behaviour from turning angles (Garriga, Palmer, Oltra, & Bartumeus, 2016; Munden et al., 2018; Potts et al., 2018), (iii) multi-second tri-axial acceleration and bearing (Bidder et al., 2015; Willener, Handrich, Halsey, & Strike, 2016; Hernández-Pliego, Rodríguez, Dell’Omo, & Bustamante, 2017; Williams et al., 2017), and/or (iv) validation datasets for supervised machine learning (Resheff, Rotics, Harel, Spiegel, & Nathan, 2014; Leos-Barajas et al., 2017). Light-level geolocators however (i) cannot provide spatially accurate enough information to infer turning angles. Furthermore, due to the weight restrictions of using them on small aerial species, they (ii) can only collect data over minutes or hours (not milliseconds). Finally, (iii) they are most commonly deployed on migratory animals that are physically impossible to follow, making training datasets impossible to collect for supervised machine learning.

Behavioural analyses of multi-sensor geolocator data therefore differ fundamentally from any previously developed behavioural classification methods for GPS tags, because behaviour must be identified independent of location. Here, we introduce **pamlr,** a toolbox for analysing behaviour from light/level geolocators using **P**ressure, **A**cceleration, temperature, **M**agnetism and **L**ight data in **R** (R Core Team, 2019). The package combines functions (Figure 1) for importing data from different multi-sensor geolocator tags (SOI-GDL3pam), functions for calculating, projecting and plotting data, and wrappers for different classification algorithms (clustering algorithms and hidden markov models) to infer behaviour. We also introduce functions specifically developed for identifying migratory flight bouts, and for collecting summary statistics from these tags. Parameters are pre-set to facilitate pattern recognition and therefore behavioural classification in birds, but is flexible and can be used for other species. Finally, *pamlr* includes functions for comparing the agreement between different model outputs. Full code can be accessed at https://kiranlda.github.io/PAMLrManual/index.html.

**Figure 1.**
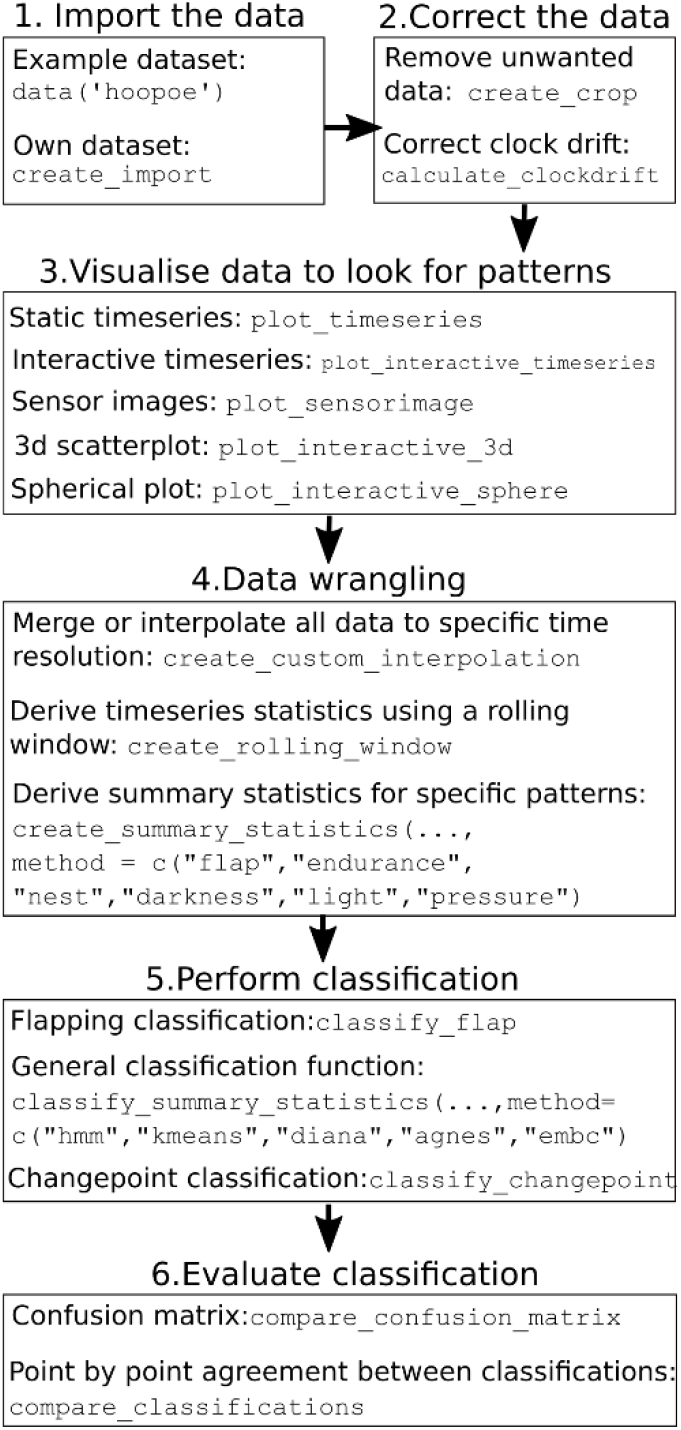
Package workflow

## 2 Tag data overview

For all loggers, data intervals and sensors are logger-dependent and customisable, requiring trade-offs between logging frequency, weight and battery life. Note that we do not describe depth-time pressure (DTP) recorders used for tracking diving marine creatures because the methods have been previously been presented (Tremblay, Cherel, Oremus, Tveraa, & Chastel, 2003). We instead focus on smaller devices with different sensors developed for tracking aerial creatures such as birds and bats (Brlík et al., 2020; Voigt, Kravchenko, Liechti, & Bumrungsri, 2020). Indeed, ***atmospheric pressure*** can be used to calculate altitude, and *pamlr* includes a function `calculate_altitude()` for this purpose (International Organization for Standardization, 1975). Many tags also record ***temperature***. Some have two sensors, one under and another on top of the device, to capture the temperature of the animal and the temperature of the atmosphere. However, body heat and feathers can bias measurements. ***Acceleration*** data is also often recorded in three formats: raw ***tri-axial acceleration*** data, ***activity***, and ***pitch***. Both activity and pitch are summary statistics calculated from the raw tri-axial data to save memory. The function `calculate_triaxial_accelerometer()` calculates roll, pitch and yaw from this tri-axial acceleration data (Bidder et al., 2015). Finally, ***magnetic field*** data is recorded on devices which also record tri-axial acceleration. Because magnetic data recording can be distorted by the presence of ferrous materials or magnetism near the sensor, the function `calculate_triaxial_magnetic()` was developed following calibration methods of Bidder et al., (2015).

Here, we illustrate the use of ***pamlr*** using SOI-GDL3pam logger data. However, the functions described in this manuscript are also applicable to other multi-sensor light loggers if formatted in a similar manner. We therefore encourage anyone using multisensor geolocator data which cannot be read by pamlr to contact us so that we may accommodate different data inputs.

## 3 Data import and wrangling: create_… functions

SOI-GDL3pam data can be imported into R using the function `create_import()`. Importantly, many tags often start recording before and after they are attached to an animal, and therefore require these periods to be cropped from the dataset using the function `create_crop()`. Furthermore, data from different sensors are collected at different temporal resolutions, `create_custom_interpolation()` formats data to the same time intervals as a specified variable (e.g. pressure) and has options to summarise finer resolution data (median, sum or snapshot) and interpolating (if desired) lower resolution data. However, interpolation is not always advisable, and another alternative for formatting data for analysis is to use a rolling window with `create_rolling_window()`, which progresses across all the timeseries and creates summary statistics for the data contained within that window of a certain timeframe, and creates a summary (minimum, maximum, or mean).

## 4 Visualisation: plot_… functions

***Time series*** are a commonly used method of plotting biologging data (Figure 2a) and pamlr offers summary plots of multiple sensors with `plot_timeseries()`. However, such plots can become noisy with large datasets, and *pamlr* offers interactive timeseries plotting for synchronising zooms across timeseries to look for patterns across sensors using `plot_interactive_timeseries()`(Vanderkam, Allaire, Owen, Gromer, & Thieurmel, 2018). To compare different datasets, the function `plot_sensorimage()` plots biologging data as an image (hereafter ***sensor image***; Figure 2b). ***Histograms*** and ***3D plots*** can also help with data interpretation by visualising data aggregation or partitioning, using the functions `plot_histogram()` and `plot_interactive_3d()` respectively (Figure 2c and d). Finally, tri-axial magnetic bearing and acceleration can be plotted onto an ***m-sphere*** (Williams et al., 2017) or ***g-sphere*** (Wilson et al., 2016), using the function `plot_interactive_sphere()` (Figure 2e).

**Figure 2.**
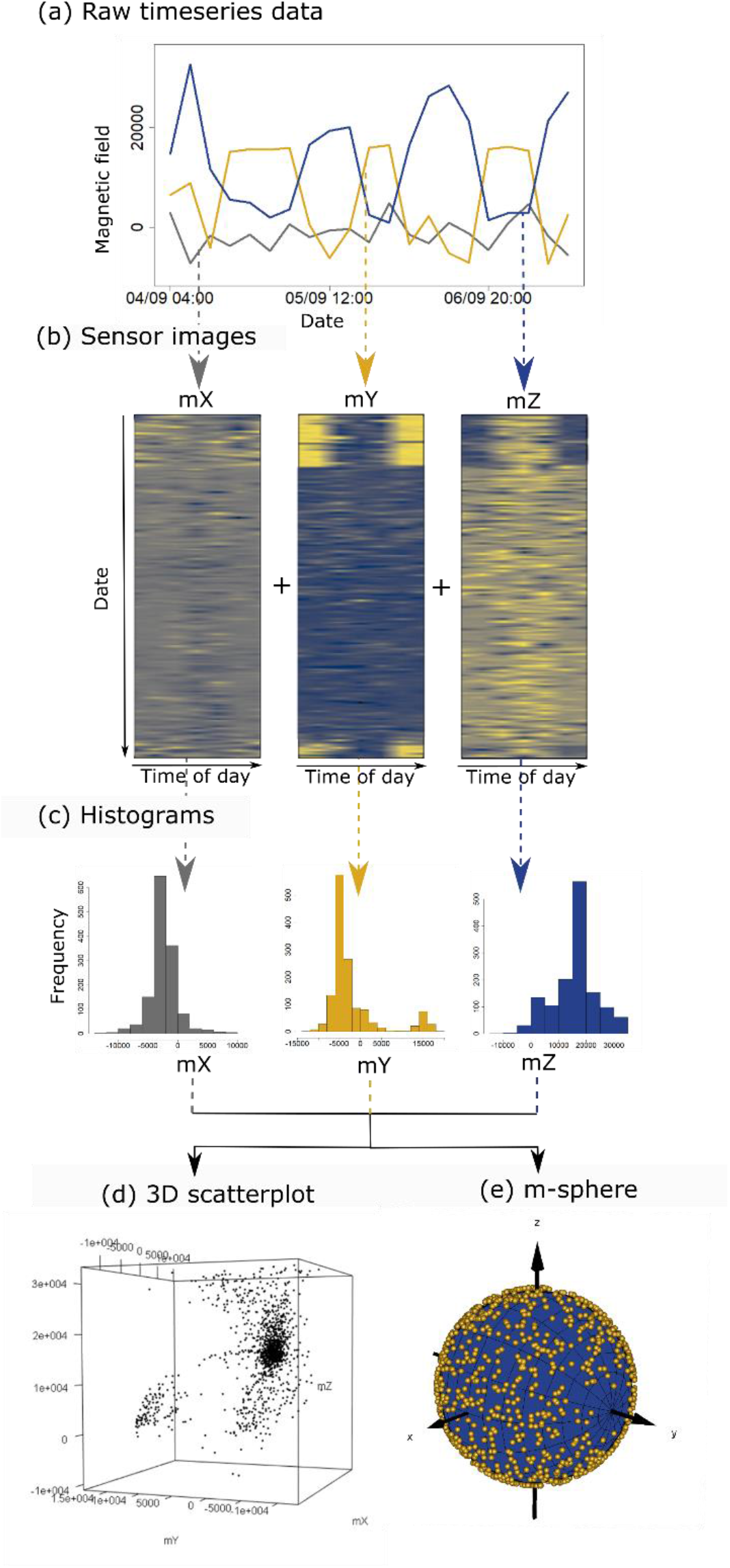
Different visualisations of magnetic field data for alpine swift *Tachymarptis melba*. To gain an initial impression of the (a) raw data, it can fist be plotted as an interactive time series. However, a great deal of insight can also be gleaned from plotting the data as (b) a sensor image. These suggest that resting periods should be easy to distinguish from others using mY as confirmed by (c) histograms and (d) 3D plots. Data can also be visualised without distortions with (e) an m-sphere.

## 5 Classifying behaviour

Data patterns can vary substantially from species to species (Figure 3). It is therefore important to think about the ecology of the species to determine what patterns might answer the question that is being asked. Functions are therefore setup to be applicable across species, however there are some predefined functions

**Figure 3.**
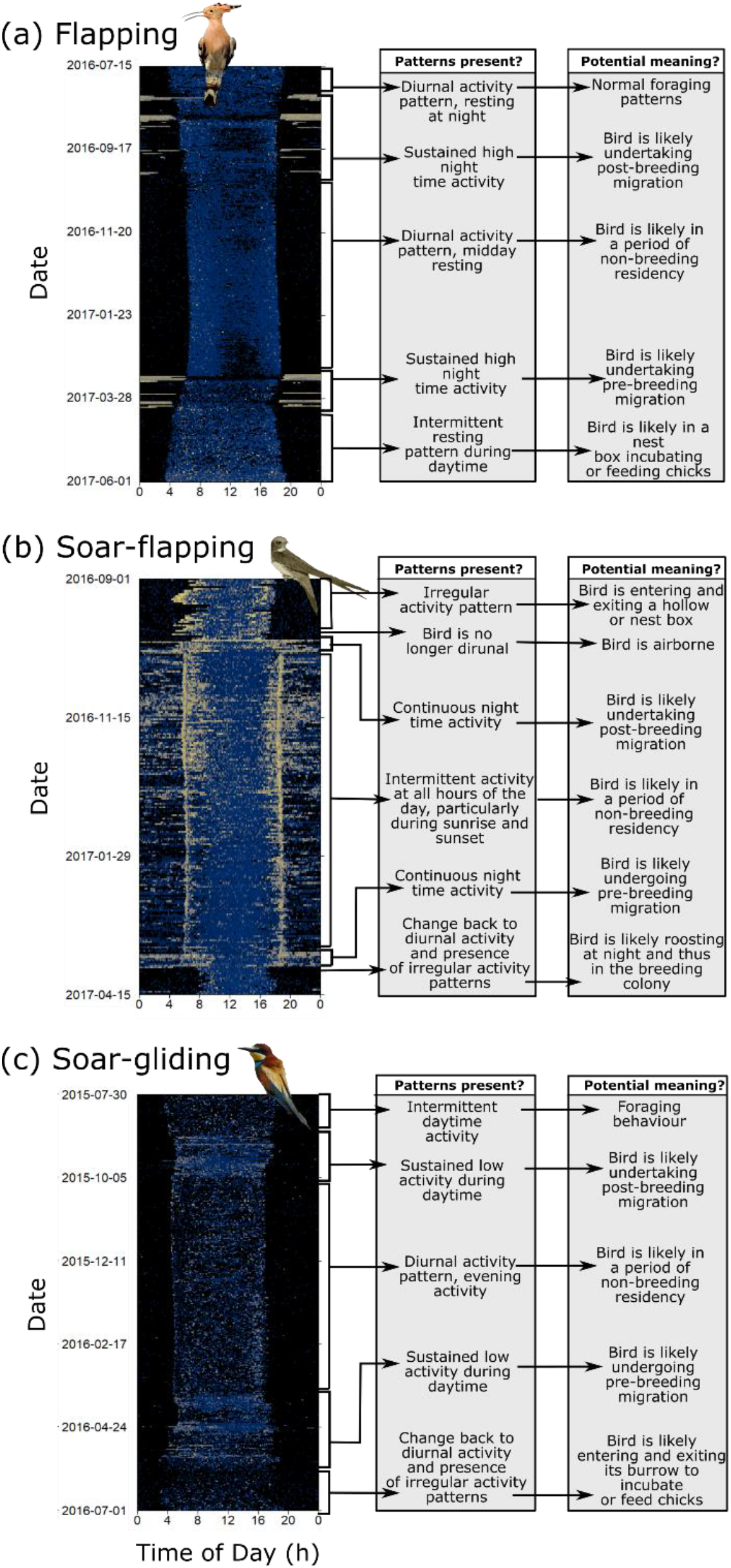
Data patterns differ substantially from species to species. Here we illustrate this first step by using actograms (left column). However, note that plotting multiple sensor images side by side can provide much richer information. In panel (a), we describe for a flapping bird (hoopoe *Epupa epops*) the activity patterns (middle column) and the behaviours they likely represent (right column). In panel (b) we do the same for a flap-gliding bird (alpine swift *Tachymarptis melba*) and in panel (c) a soar-gliding bird (European bee-eater *Merops apiaster*).

### 5.1 Classifying migratory flight bouts in passerines

One of the most common and useful applications of pamlr, is the estimation of migratory flight duration. Compared to packages such as `GeoLight` that use variations in daylight hours to calculate migratory timetables with the function `changeLight` (Simeon Lisovski & Hahn, 2012; S. Lisovski et al., 2020), pamlr instead uses the activity sensor to identify migratory flight bouts in birds with flapping flight behaviour in the function `classify_flap()` (Figure 4). The estimated migratory timetable is therefore much more precise than that estimated from light alone (as seen in Bäckman et al., 2017; Liechti et al., 2018; Sjöberg et al., 2018). The function outputs a migratory timetable.

**Figure 4.**
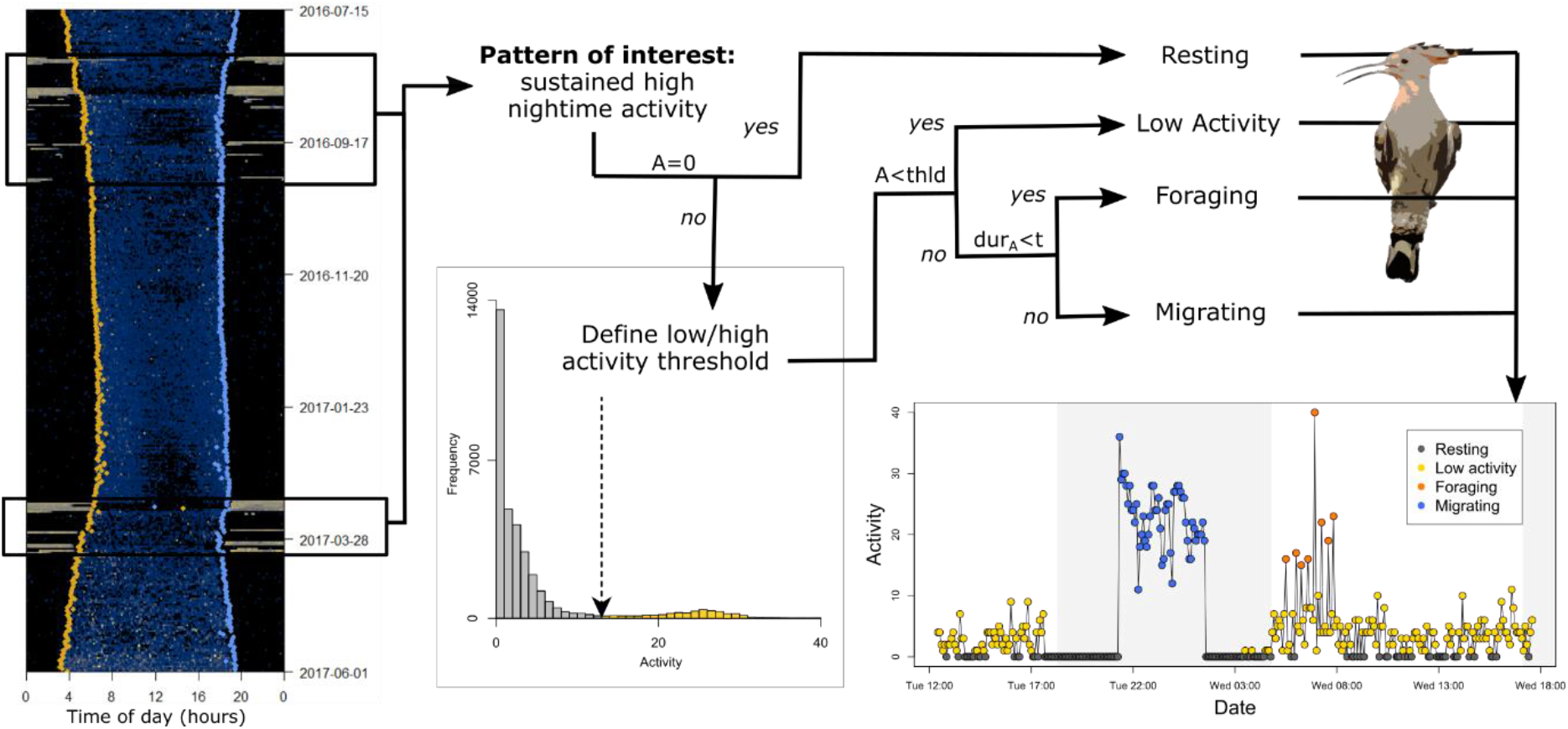
Schematic representation of the `classify_flap()` algorithm for classifying flapping migratory behaviour.

### 5.2 Common data patterns for birds

There are a number of other behaviours that are identifiable using multisensory geolocator tags. Though these tags have primarily been used on birds (Bäckman et al., 2017; Dhanjal-Adams et al., 2018; Liechti et al., 2018; Sjöberg et al., 2018; Briedis, Beran, Adamík, & Hahn, 2020; Evens et al., 2020; Barras, Arlettaz, & Liechti, 2021; Sander et al., 2021), they can also be used with bats (Voigt et al., 2020). There are therefore a number of behaviours that might be analysed from these tags. The function `create_summary_statistics()` is setup to allow the user to extract particular patterns from the raw data that may be useful to classify particular behaviours using some of the methods define in section 5.3.

Depending on the behaviour (see examples in Figure 3), flight bouts can be characterised by (i) continuous high activity which can be extracted from the data using the method `“flap”`, (ii) sustained activity (low and high) using `“endurance”`, (iii) a pressure change greater than the background pressure changes due to weather using `“pressure”`, (iv) a period of continuous light using `“light”`. Incubation bouts can be characterised by (v) periods of darkness using `“darkness”` and finally (vi) periods of resting using `“rest”`. These functions also calculate summary statistics for each event. These include, but are not limited to, how much the animal changed altitude during the event, how active it was during that event, whether it was night or day during that event, how long the event lasted, how many other similar events occurring during the same day, how often these events lasted overall that day, whether pressure at the start of the event was different from pressure at the end.

### 5.3 Classification methods: classify_… functions

Data created by the function `create_summary_statistics()` can then be classified to extract particular behaviours. Indeed, pamlr offers options for classifying data using a ***changepoint analysis*** to delineate differences in variance, in mean or in both. The default setting in `classify_changepoint()` are set for finding differences between migratory and non-migratory periods using pressure, but the user can customise these to fit their needs (see Killick, Haynes, Eckley, Fearnhead, & Lee, 2016).

More detailed multiclass classifications can also be implemented using `classify_summary_statistics()` for cluster analysis and hidden Markov models. Indeed, pamlr integrates three types of clustering algorithms: *k*-means, hierarchical and expectation-minimisation binary clustering. ***k-means clustering*** minimises the within-cluster sum of squares of the points (Hartigan & Wong, 1979) and can be implemented by using the method `“kmeans”`. ***Hierarchical clustering*** analyses can either be agglomerative or divisive (see Maechler et al., 2018). The first relies on putting each datapoint in a single cluster and successively merging them until a stopping criterion is satisfied and can be implemented using `“agnes”`. The divisive hierarchical clustering method starts by allocating all datapoints to the same cluster and splitting each cluster until a stopping criterion is met. This can be implemented using `“diana”`. ***Expectation-minimisation binary clustering*** (EMbC), clusters data based on geometry alone (Garriga et al., 2016). More specifically binary delimiters are used to segregate the data along an axis, forcing centroids to lie within these binary regions. Analysis can be undertaken using `“embc”`.

***Hidden Markov Models*** (HMMs) are stochastic time-series models (see Visser & Speekenbrink, 2010) which assume that the observed time series (such as the measured acceleration, temperature or pressure) is driven by an unobservable state process (such as flying or resting behaviour) and can be implemented using `“hmm”`.

## 6 Measuring classification accuracy: compare_… functions

Not all methods should be used exclusively. Some can be used to complement each other, while others can be used interchangeably. pamlr offers a function `compare_classification()` which takes multiple classification outputs and summarises the agreement between all of these. The function `compare_confusion_matrix()` also populates a confusion matrix using predicted and reference points. If no reference data are available, two different classifications can instead be compared.

## 7 Outlook

Here, we present functions adapted to the analysis of multisensor geolocator tags with particular focus on bird behaviour. Indeed, the classifications of migratory flight periods can for instance be used to increase the precision of geolocation estimates (Simeon Lisovski et al., n.d.) and can answer a suite of biological questions. Importantly, many of the functions in pamlr are setup to be generalisable and applicable to animals other than birds. Miniaturised activity, pitch, pressure and temperature sensors can provide important information on the natural history, behaviour and physiology of any species too small to carry a GPS. Furthermore, many multi-sensor geolocator tags are now customisable. Thus, the temporal data resolution of, for instance, tri-axial accelerometer and magnetometer recordings on SOI-GDL3pam loggers could be increased with a shorter battery life, allowing more detailed and complex behavioural classifications to be performed over smaller time scales, such as dead-reckoning (Bidder et al., 2015). From these reconstructed tracks and turning angles, many previously developed finer resolution analyses can also be applied (e.g. Garriga et al., 2016; Potts et al., 2018). The collection of observation data would also allow for the development of supervised machine learning methods (Valletta, Torney, Kings, Thornton, & Madden, 2017). Multisensor geolocator tags therefore provide exciting new opportunities for analysing otherwise unseen behaviours in animals that were previously impossible to tag.

## 8 Author contributions

KLDA led the writing of both package and manuscript. All authors contributed towards the conceptualisation of the methods, and towards editing the manuscript.

## 9 Acknowledgements

The Swiss federal office for environment contributed financial support for the development of the data loggers (UTF-Nr. 254, 332, 363, 400).

## References

Bäckman, J., Andersson, A., Alerstam, T., Pedersen, L., Sjöberg, S., Thorup, K., & Tøttrup, A. P. (2017). Activity and migratory flights of individual free-flying songbirds throughout the annual cycle: method and first case study. Journal of Avian Biology, 48(2), 309–319. doi: 10.1111/jav.01068

Barras, A. G., Arlettaz, R., & Liechti, F. (2021). Complex Day-to-day Movements of an Alpine Passerine May Act as an Insurance Against Environmental Variability. doi: 10.21203/rs.3.rs-184098/v2

Bidder, O. R., Walker, J. S., Jones, M. W., Holton, M. D., Urge, P., Scantlebury, D. M., … Wilson, R. P. (2015). Step by step: reconstruction of terrestrial animal movement paths by dead-reckoning. Movement Ecology, 3(1), 23. doi:10.1186/s40462-015-0055-4

Briedis, M., Beran, V., Adamík, P., & Hahn, S. (2020). Integrating light-level geolocation with activity tracking reveals unexpected nocturnal migration patterns of the tawny pipit. Journal of Avian Biology, 51(9), jav.02546. doi:10.1111/jav.02546

Brlík, V., Koleček, J., Burgess, M., Hahn, S., Humple, D., Krist, M., … Procházka, P. (2020). Weak effects of geolocators on small birds: A meta-analysis controlled for phylogeny and publication bias. Journal of Animal Ecology, 89(1), 207–220. doi:10.1111/1365-2656.12962

Dhanjal-Adams, K. L., Bauer, S., Emmenegger, T., Hahn, S., Lisovski, S., & Liechti, F. (2018). Spatiotemporal Group Dynamics in a Long-Distance Migratory Bird. Current Biology, 28(17), 2824–2830.e3. doi:10.1016/j.cub.2018.06.054

Dreelin, R. A., Shipley, J. R., & Winkler, D. W. (2018). Flight Behavior of Individual Aerial Insectivores Revealed by Novel Altitudinal Dataloggers. Frontiers in Ecology and Evolution, 6, 182. doi:10.3389/fevo.2018.00182

Edwards, E. W. J., Quinn, L. R., & Thompson, P. M. (2016). State-space modelling of geolocation data reveals sex differences in the use of management areas by breeding northern fulmars. Journal of Applied Ecology, 53(6), 1880–1889. doi:10.1111/1365-2664.12751

Evens, R., Kowalczyk, C., Norevik, G., Ulenaers, E., Davaasuren, B., Bayargur, S., … Kempenaers, B. (2020). Lunar synchronization of daily activity patterns in a crepuscular avian insectivore. Ecology and Evolution, 10(14), 7106–7116. doi:10.1002/ece3.6412

Frisius, R. G. (1544). De principiis astronomiae & cosmographiae deque usu blobi ab eodem editi. in aedibus Ioan. Stelsii, 1548. Retrieved from https://books.google.ch/books?id=IEFmAAAAcAAJ

Garriga, J., Palmer, J. R. B., Oltra, A., & Bartumeus, F. (2016). Expectation-Maximization Binary Clustering for Behavioural Annotation. PLOS ONE, 11(3), e0151984. doi: 10.1371/journal.pone.0151984

Hartigan, J. A., & Wong, M. A. (1979). A K-Means Clustering Algorithm. Applied Statistics, 28(1), 100. doi:10.2307/2346830

Hedenström, A., Norevik, G., Warfvinge, K., Andersson, A., Bäckman, J., & Åkesson, S. (2016). Annual 10-Month Aerial Life Phase in the Common Swift Apus apus. Current Biology, 26(22), 3066–3070. doi:10.1016/J.CUB.2016.09.014

Hernández-Pliego, J., Rodríguez, C., Dell’Omo, G., & Bustamante, J. (2017). Combined use of tri-axial accelerometers and GPS reveals the flexible foraging strategy of a bird in relation to weather conditions. PLOS ONE, 12(6), e0177892. doi:10.1371/journal.pone.0177892

International Organization for Standardization. (1975). International organization for standardization. ISO, 2533, 1975.

Killick, R., Haynes, K., Eckley, I., Fearnhead, P., & Lee, J. (2016). Package ‘changepoint’ Type Package: Methods for Changepoint Detection. CRAN. Retrieved from https://cran.r-project.org/web/packages/changepoint/changepoint.pdf

Leos-Barajas, V., Photopoulou, T., Langrock, R., Patterson, T. A., Watanabe, Y. Y., Murgatroyd, M., & Papastamatiou, Y. P. (2017). Analysis of animal accelerometer data using hidden Markov models. Methods in Ecology and Evolution, 8(2), 161–173. doi:10.1111/2041-210X.12657

Liechti, F., Bauer, S., Dhanjal-Adams, K. L., Emmenegger, T., Zehtindjiev, P., & Hahn, S. (2018). Miniaturized multi-sensor loggers provide new insight into year-round flight behaviour of small trans-Sahara avian migrants. Movement Ecology, 6(1), 19. doi:10.1186/s40462-018-0137-1

Liechti, F., Witvliet, W., Weber, R., & Bächler, E. (2013). First evidence of a 200-day non-stop flight in a bird, 4, 2554. Retrieved from http://dx.doi.org/10.1038/ncomms3554

Lisovski, S., Bauer, S., Briedis, M., Davidson, S. C., Dhanjal-Adams, K. L., Hallworth, M. T., … Bridge, E. S. (2020). Light-level geolocator analyses: A user’s guide. Journal of Animal Ecology, 89(1). doi:10.1111/1365-2656.13036

Lisovski, Simeon, Bauer, S., Briedis, M., Davidson, S., Dhanjal-Adams, K., Karagicheva, J., … Bridge, E. (n.d.). Light-Level Geolocator Analyses: A user’s guide. Journal of Animal Ecology.

Lisovski, Simeon, & Hahn, S. (2012). GeoLight–processing and analysing light-based geolocator data in R. Methods in Ecology and Evolution, 3(6), 1055–1059.

Lisovski, Simeon, Hewson, C. M., Klaassen, R. H. G., Korner-Nievergelt, F., Kristensen, M. W., & Hahn, S. (2012). Geolocation by light: accuracy and precision affected by environmental factors. Methods in Ecology and Evolution, 3(3), 603–612. doi:10.1111/j.2041-210X.2012.00185.x

Maechler, M., Rousseeuw, P., Struyf, A., Hubert, M., Hornik, K., Studer, M., … Kozlowski, K. (2018). Package ‘cluster’: Finding Groups in Data. CRAN. Retrieved from https://cran.r-project.org/web/packages/cluster/cluster.pdf

Meier, C. M., Karaardıç, H., Aymí, R., Peev, S. G., Bächler, E., Weber, R., … Liechti, F. (2018). What makes Alpine swift ascend at twilight? Novel geolocators reveal year-round flight behaviour. Behavioral Ecology and Sociobiology, 72(3), 45. doi:10.1007/s00265-017-2438-6

Munden, R., Börger, L., Wilson, R. P., Redcliffe, J., Loison, A., Garel, M., & Potts, J. R. (2018). Making sense of ultrahigh-resolution movement data: A new algorithm for inferring sites of interest. Ecology and Evolution, 9(1), 265–274. doi:10.1002/ece3.4721

Potts, J. R., Börger, L., Scantlebury, D. M., Bennett, N. C., Alagaili, A., & Wilson, R. P. (2018). Finding turning-points in ultra-high-resolution animal movement data. Methods in Ecology and Evolution, 9(10), 2091–2101. doi:10.1111/2041-210X.13056

R Core Team. (2019). R: A Language and Environment for Statistical Computing. Vienna, Austria: Foundation for Statistical Computing. Retrieved from https://www.r-project.org/

Resheff, Y. S., Rotics, S., Harel, R., Spiegel, O., & Nathan, R. (2014). AcceleRater: a web application for supervised learning of behavioral modes from acceleration measurements. Movement Ecology, 2(1), 27. doi: 10.1186/s40462-014-0027-0

Sander, M., Chamberlain, D., Alba, R., Jähnig, S., Rosselli, D., & Lisovski, S. (2021). Reduced Breeding Success Suggests Trophic Mismatch Despite Timely Arrival in an Alpine Songbird. doi:10.21203/rs.3.rs-137126/v1

Shaffer, S. A., Tremblay, Y., Awkerman, J. A., Henry, R. W., Teo, S. L. H., Anderson, D. J., … Costa, D. P. (2005). Comparison of light- and SST-based geolocation with satellite telemetry in free-ranging albatrosses. Marine Biology, 147(4), 833–843. doi:10.1007/s00227-005-1631-8

Sjöberg, S., Pedersen, L., Malmiga, G., Alerstam, T., Hansson, B., Hasselquist, D., … Bäckman, J. (2018). Barometer logging reveals new dimensions of individual songbird migration. Journal of Avian Biology, 49(9), e01821. doi:10.1111/jav.01821

Tremblay, Y., Cherel, Y., Oremus, M., Tveraa, T., & Chastel, O. (2003). Unconventional ventral attachment of time–depth recorders as a new method for investigating time budget and diving behaviour of seabirds. The Journal of Experimental Biology, 206, 1929–1940. doi: 10.1242/jeb.00363

Valletta, J. J., Torney, C., Kings, M., Thornton, A., & Madden, J. (2017). Applications of machine learning in animal behaviour studies. Animal Behaviour, 124, 203–220. doi:10.1016/J.ANBEHAV.2016.12.005

Vanderkam, D., Allaire, J., Owen, J., Gromer, D., & Thieurmel, B. (2018). ‘Dygraphs’ Interactive Time Series Charting Library. Dygraphs: Interface R. Retrieved from https://cran.r-project.org/package=dygraphs

Visser, I., & Speekenbrink, M. (2010). depmixS4: an R package for hidden Markov models. Journal of Statistical Software, 36(7), 1–21.

Voigt, C. C., Kravchenko, K., Liechti, F., & Bumrungsri, S. (2020). Skyrocketing Flights as a Previously Unrecognized Behaviour of Open-Space Foraging Bats. Acta Chiropterologica, 21(2), 331–339. doi:10.3161/15081109ACC2019.21.2.008

Willener, A. S. T., Handrich, Y., Halsey, L. G., & Strike, S. (2016). Fat King Penguins Are Less Steady on Their Feet. PLOS ONE, 11(2), e0147784. doi:10.1371/journal.pone.0147784

Williams, H. J., Holton, M. D., Shepard, E. L. C., Largey, N., Norman, B., Ryan, P. G., … Wilson, R. P. (2017). Identification of animal movement patterns using tri-axial magnetometry. Movement Ecology, 5(1), 6. doi:10.1186/s40462-017-0097-x

Wilson, R. P., Holton, M. D., Walker, J. S., Shepard, E. L. C., Scantlebury, D. M., Wilson, V. L., … Jones, M. W. (2016). A spherical-plot solution to linking acceleration metrics with animal performance, state, behaviour and lifestyle. Movement Ecology, 4(1), 22. doi: 10.1186/s40462-016-0088-3

